# High-resolution cryo-EM reconstructions in the presence of substantial aberrations

**DOI:** 10.1101/798280

**Authors:** Raquel Bromberg, Yirui Guo, Dominika Borek, Zbyszek Otwinowski

**Author notes:** corresponding authors Dominika Borek, Department of Biophysics, The University of Texas Southwestern Medical Center, Dallas, Texas 75390, USA, Zbyszek Otwinowski, Department of Biophysics, The University of Texas Southwestern Medical Center, Dallas, Texas 75390, USA.

## Abstract

The beam-image shift method accelerates data acquisition in cryo-EM single particle reconstruction (cryo-EM SPR) by fast repositioning of the imaging area, but at the cost of more severe and complex optical aberrations.

We analyze here how uncorrected anti-symmetric aberrations, such as coma and trefoil, affect cryo-EM SPR results, and then infer an analytical formula quantifying information loss due to their presence that explains why Fourier-shell coefficient (FSC)-based statistics may report significantly overestimated resolution if these aberrations are not fully corrected. We validate our analysis with reference-based aberration refinement for two cryo-EM SPR datasets acquired with a 200 kV microscope in the presence of coma exceeding 40 µm, and obtained 2.3 and 2.7 Å reconstructions for 144 and 173 kDa particles, respectively.

Our results provide a description of an efficient approach for assessing information loss in cryo-EM SPR data acquired in the presence of higher-order aberrations and address inconsistent guidelines regarding the level of aberrations acceptable in cryo-EM SPR experiments.

## Introduction

In a typical cryo-EM SPR experiment, some aberrations such as defocus are introduced intentionally while others such as spherical aberration are unavoidable for a given setup (1-3). The significance of the remaining aberrations is evaluated on a case-by-case basis (4, 5).

In the analysis of aberrations in cryo-EM SPR, where in one image we record only a small part of the focal plane, we can use an isoplanatic approximation in which aberrations are represented by convolutions, and in Fourier space they depend only on the angle of the scattered electrons. In phase contrast illumination mode, used in cryo-EM SPR, axial aberrations can be divided into two distinct categories depending on the symmetry properties of the image phase shift as a function of a scattering vector. If the phase shift is centrosymmetric, then aberrations will result in modulations of the image power spectrum. If the phase shift is antisymmetric, the power spectrum will not be modulated because aberrations will not affect the amplitude of the image but only its phase. The lowest order of antisymmetric phase shift is a translation, which due to the lack of an absolute coordinate system, can be set to zero. Antisymmetric aberrations of the next, third order are called axial coma and trefoil and these are important for cryo-EM SPR data quality in practice (4-6). Alignment procedures used in cryo-EM SPR minimize coma and trefoil indirectly, for instance by analyzing changes in the image power spectrum due to interactions between beam tilt and spherical aberration (5). The success of this approach and similar ones requires co-alignment of the optical axes for multiple lenses, and if the alignment procedures are not properly executed, coma and trefoil may be present and affect the quality of the SPR results, and yet manifest only in specialized analyses (5-8). Furthermore, these and additional, higher-order, antisymmetric aberrations are induced when data are acquired with the beam-image shift method (6-8), in which coordinated electronic shifts of an illuminating beam and an image are used to navigate away from the optical axis. We restrict our discussion here to axial aberrations, but with the understanding that these aberrations will have different values in different positions of an optical system for data acquired with the beam-image shift method.

The acceleration in cryo-EM data acquisition enabled by the beam-image shift method (6-9) has resulted in discussions of the best experimental strategies for data collection, approaches to compensate for or to correct for optical aberrations, and methods to assess modulation and the loss of signal due to the presence of uncorrected optical aberrations (4, 6, 10, 11). Axial coma can be corrected by applying compensating beam tilt during data collection; however, beam tilting does not correct other aberrations (4), and thus the extent to which one can apply beam-image shift without compromising data quality remains open.

We found that large values of axial aberrations can be precisely estimated and accurately corrected for, leading to large, case-specific improvements of SPR results. We provide formula for assessing how the levels of uncorrected coma and trefoil affect the resolution of SPR and discuss their impact on the validation statistics.

## Results

CTF determination by power spectrum analysis is an inherent part of high-resolution cryo-EM SPR and provides estimates of the magnitude of symmetric aberrations (2). However, phase shift does not modulate the power spectrum, so determination of antisymmetric aberrations has to be performed by other methods (4).

In our analysis of antisymmetric aberrations, we utilize an associative property of pure phase shift that also holds when it is combined with image translation (12). Thus, corrections for the presence of antisymmetric aberrations can be split into separate steps and applied in any order, without loss of accuracy in retrieving information. Consequently, for both large and small magnitudes of antisymmetric aberrations, their impact is defined only by the difference between their true and their assumed or refined values. If their estimates are inaccurate, the difference generates a component of the image phase shift that, after averaging over multiple particles, produces the signal modulation that we analyze here.

Antisymmetric aberration is a convolution with signal so in the frequency domain the convolution is represented as the multiplication of a signal and Fourier representation of an aberration. The impact of aberrations on 3D cryo-EM SPR has been theoretically analyzed by considering how images are affected (5, 13). However, cryo-EM SPR relies on averaging multiple particles to increase the SNR. Thus, we investigated how an aberration’s impact will propagate to averaged representations of particles.

To this end, we assume a large number of particles which are randomly oriented on a grid with respect to rotation around the beam axis but with potential preferred orientation dependence on the other two Eulerian angles. Averaging all such particles removes the dependence of the average signal on the angle of the aberration in the microscope frame (6). Therefore, the final consequence of an aberration is the resolution-dependent modulation of signal amplitude (6). For each unique projection that is conceptually equivalent to a 2D class average, we can define an angle of particle orientation φ in the microscope and the difference Δ*φ* between *φ* and the characteristic direction of a particular antisymmetric aberration (Fig. S1). Single particles have no force aligning them with this angle, so we can assume that their distribution is uniform with respect to this angle. We then express the maximum phase shift *a* for a particular aberration and resolution, which for coma and trefoil, is:

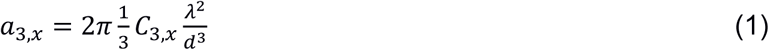

where *d* represents resolution and *λ* represents electron wavelength. The first index of the aberration coefficients identifies the power dependence on resolution and the other defines angular periodicity in the microscope frame of the phase shift, and so for coma *x* = 1 and for trefoil *x* = 3 (14). Coma can also be derived from interactions between beam tilt *b* and spherical aberrations 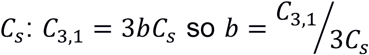. This representation may be convenient when one is interested in expressing coma with respect to beam tilt rather than coma values.

In weak phase approximation, third order aberrations can be expressed using terms resulting from the Taylor expansion of a wave front: 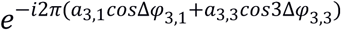. To obtain a modulation term for the signal at a given resolution, we average the wave front over the angular distribution of all possible particles, with the result being the Bessel function of order zero *J*_o_ (Fig. 1):

**Figure 1.**
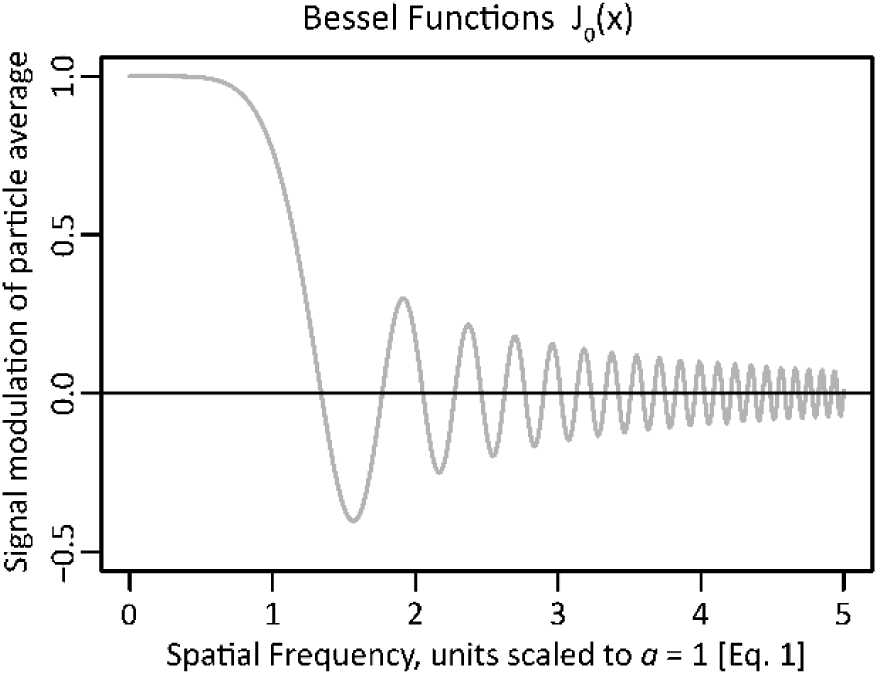
Resolution dependence of third order aberrations on single particle reconstruction. X-axis represents reciprocal space resolution, scaled to *a = 1* for *x = 1* (Eq. 1). Y-axis represents reciprocal space signal modulation of the reconstruction resulting from averaging the aberration.

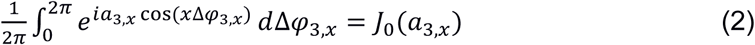

To our knowledge, this analytical result has not been noticed before in assessing aberrations despite having highly important consequences for their analysis. As discussed in detail later, the foremost consequence is that the structure factors of the reconstruction may become anticorrelated from the reality in some resolution shells, an effect which can be missed in standard, FSC-based half-maps assessment of resolution (15-17), implying much higher resolution than that achieved.

We expect that data acquired with the beam-image shift method will be affected by more than one type of axial aberration. In such a case, aberrations that have the same angular dependence order (second index) but different radial order (first index) will be strongly correlated (5) and these correlations have to be considered when values of aberrations are refined. If in refinement, the data were to have uniform information content across resolutions, the refinement will be orthogonalized by Zernike polynomials with ρ=1 corresponding to the limiting resolution (18). For coma, the corresponding Zernike polynomial is ρ^3^ − 2/3ρ with the first term representing coma and the second representing translation. Consequently, at the resolution limit, two-thirds of the coma-produced phase shifts will be compensated for by image translation in refinement. This translation can be executed on the whole image or at the level of particles by shifting their positions in then image by the same value, with such an effect not being noticeable in a typical refinement. However, the compensation factor is reduced from the value of two-thirds because the signal is much stronger at low resolution than at the resolution limit. We determined for analyzed high-resolution cryo-EM SPR datasets (Table 1) that translation produced only about a two-fifths compensating contribution, reducing the maximum value of the coma-induced phase shift by a factor of ∼0.6. The compensation due to resolution dependence explains some of the discrepancies in the literature discussing the acceptable limits of coma. For instance, one proposed limit was a 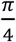 phase shift in the direction of coma distortion, and without translational compensation (4, 5). Using our formula (Eq. 2), we found that this will preserve 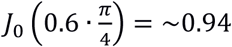 of the original signal, a reduction in the SNR that is barely significant, and so the 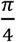 limit is too conservative as it was postulated by Cheng et al. (6). In Cheng et al.’s analysis, the impact of coma was derived from FSC curves calculated from numerical experiments, where a particular value of beam tilt (and corresponding coma) were assumed. FSC-based resolution analysis in the presence of uniform large coma can be misleading because the *J*_*0*_ function oscillates (Fig. 1). When the signal modulation term defined by *J*_*0*_ (Eq. 2) is negative, both halves of the split data are affected, and so the correlation coefficient between halves is positive even if the resulting reconstruction has a negative correlation with the truth. If the coma causes very strong modulation in the FSC (11), then it is easy to recognize the correct resolution limit corresponding to the first zero of the Bessel function *J*_*0*_ (Eq. 1, 2). However, oscillations in the FSC curve may be pronounced or smoothed to different degrees (Fig. 2), even when data sets with very similar values of coma (Table 2) were analyzed. The method that we employed from cisTEM (19) uses only a smooth spherical mask to calculate FSC curves, and so the oscillations could not have come from a molecular mask effect (20). Thus, in the absence of pronounced oscillations, the resolution limit may be significantly overestimated compared to the consideration based on the first zero of the Bessel function. Consequently, simulation-based procedures relying solely on FSC (6) may grossly underestimate the significance of coma.

**Table 1.**
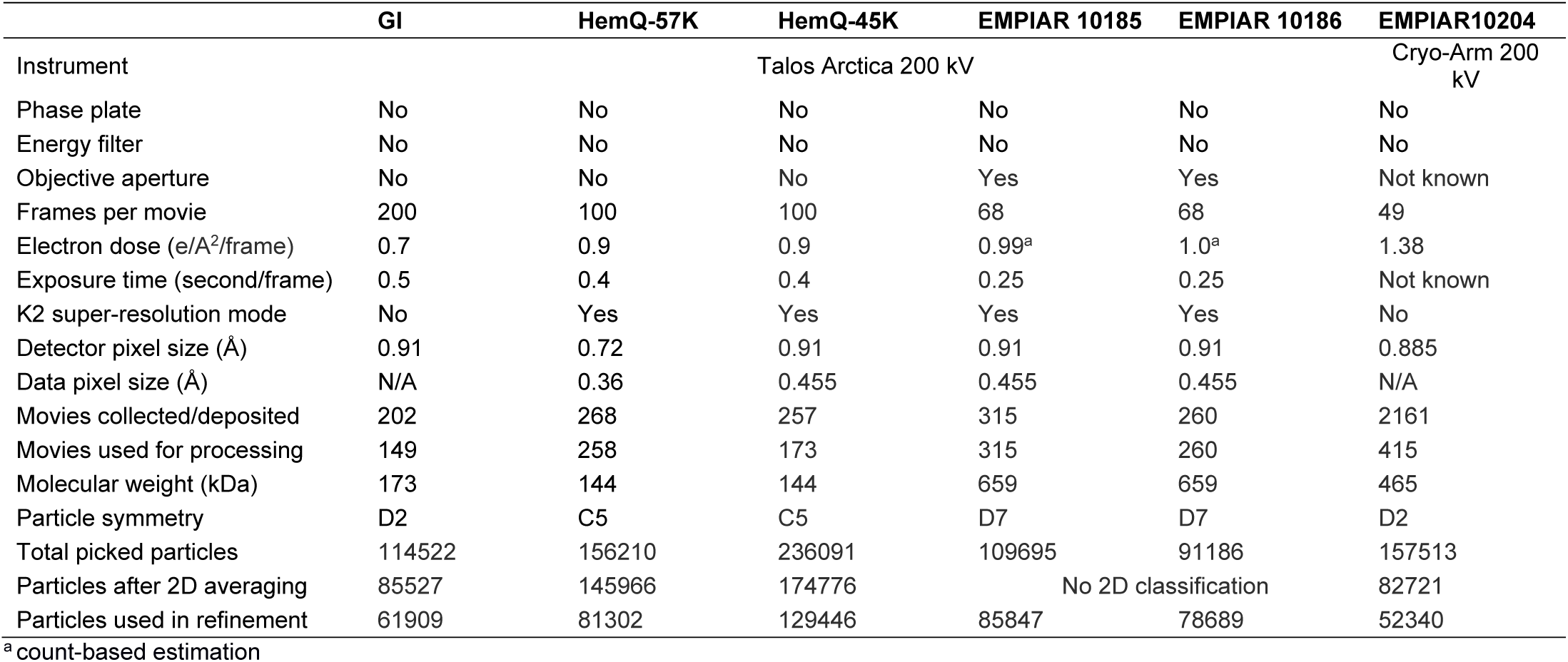
Data collection and processing

**Table 2.** The resolution without and with correction for coma for analyzed datasets.

|  | GI | HemQ-45 | HemQ-57 | EMPIAR<br>10204 | EMPIAR<br>10185 | EMPIAR<br>10186 |
| --- | --- | --- | --- | --- | --- | --- |
| Reported resolution [Å] | NA | NA | NA | NA | 3.1 | 3.3 |
| FSC<sub>0.143</sub> based resolution<br>before correction [Å] | 4.1 | 3.8 | 4.3 | 2.5 | 3.1 | 3.2 |
| FSC<sub>0.143</sub> based resolution<br>after correction [Å] | 2.7 | 2.6 | 2.3 | 2.5 | 2.5 | 2.4 |
| Coma [μm]/Beam tilt [mrad] | 42.7/5.3 | 56.9/7.0 | 56.2/6.9 | Varied<sup>a</sup> | Substantial<br>Trefoil and<br>Variable<br>coma<sup>b</sup> | Substantial<br>Trefoil and<br>Variable<br>coma<sup>b</sup> |
| Trefoil [μm] | 0.62 | 0.79 | 0.49 | 0.09 | 2.74 | 3.06 |
| Resolution at the first<br>oscillation of $J_0$ with 40%<br>compensation | 5.2 | 5.7 | 5.7 | ND | ND | ND |

<sup>a</sup>Coma varied between movies by more than 10 μm indicating ~1.2 mrad beam tilt variation
<sup>b</sup>EMPIAR 10185 and 10186 were collected consecutively and share the same stable value of trefoil. EMPIAR 10185 was collected with stage shift and has similar coma variation as EMPIAR 10204. EMPIAR 10186 was collected with beam-image shift that induced additional coma variation.

**Figure 2.**
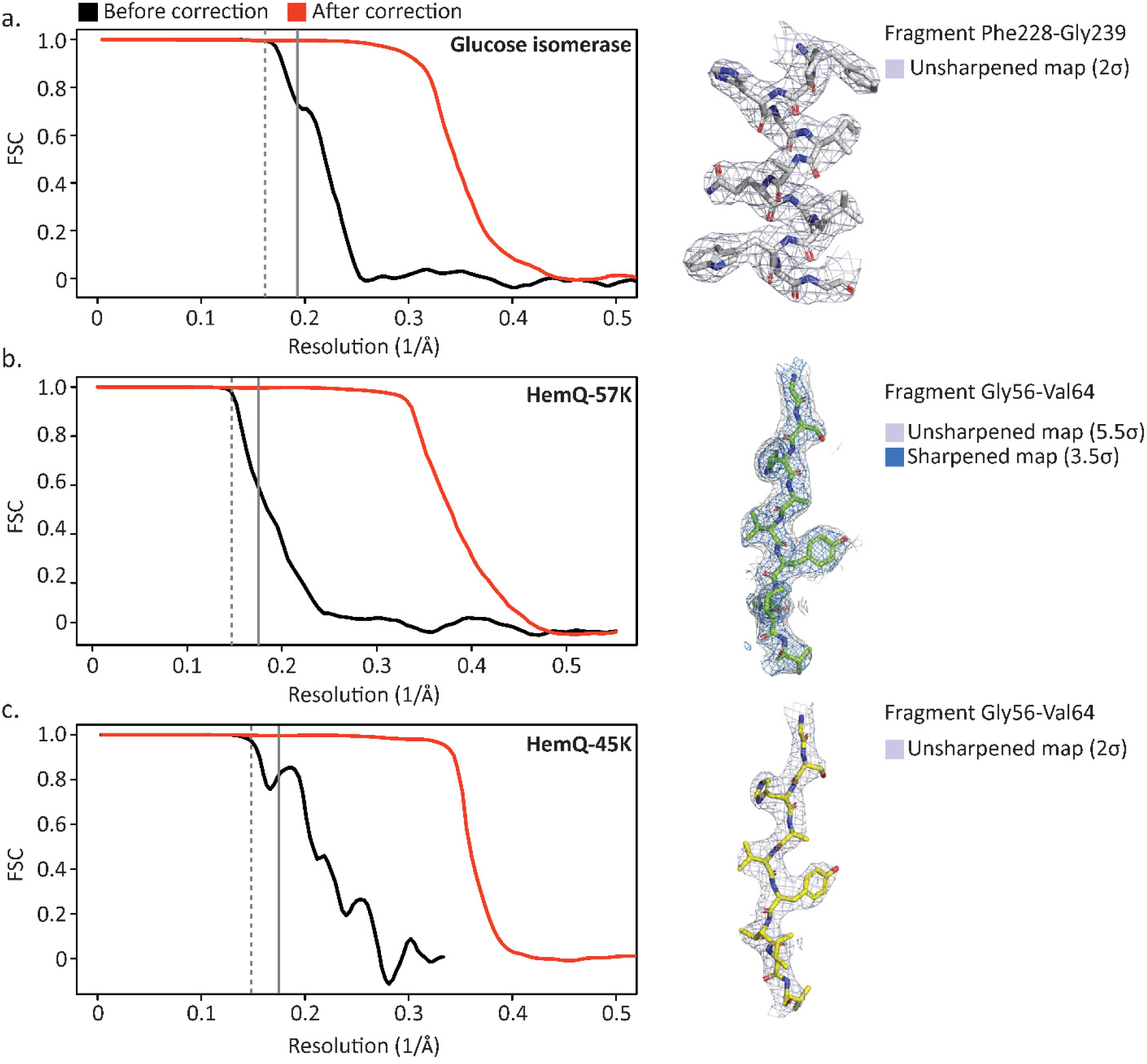
FSC plots before (black) and after (red) coma correction and the corresponding final map fragment for three experiments with high beam tilt (coma) values. The statistics from each experiment are presented in Table 1. The vertical dotted grey line represents the first zero of the modulation function (Eq. 1) and the solid line represents the first zero of the modulation function with the assumption that 40% of the coma impact was compensated by image translation. The resolutions corresponding to the first zero values are listed in Table 2.

We tested our approach for two small proteins with particle size 144 kDa and 173 kDa (Fig. 2, Table 1), and we obtained reconstructions of 2.32 Å and 2.70 Å, respectively, in the presence of very high coma, with data acquired with Talos Arctica 200 kV, K2 Gatan camera. An objective aperture, an energy filter, and a phase plate were not used, and numbers of micrographs and particles in data analysis were moderate (Table 1). To our knowledge, these are the highest resolution reconstructions obtained with a 200 kV instrument for particles having molecular mass below 200 kDa and the first high resolution reconstruction for a molecule with mass below 150 kDa. We have not found any detrimental effects for correcting coma, even in cases where the generated phase shift is very large, on the order of 10×2π (7.2 mrad). We reprocessed data from EMPIAR for 200 kV instruments, applying the same aberration estimation approach to data for the larger molecules of the proteasome (700 kDa, EMPIAR 10185 and 10186) (10) and β-galactosidase (430 kDa, EMPIAR 10204) (Table 1) (21). We refined coma and trefoil independently on each micrograph. We noticed that coma can fluctuate far above refinement uncertainty and we attribute this observation to differences in the stability of the beam tilt direction between micrographs. In addition, coma refinement has a strong correlation with overall image shift, affecting the accuracy of coma determination. We found that trefoil on the other hand was remarkably stable and has no significant correlation with other parameters of refinement, so variations in its refined value can be used as a good indicator of statistical uncertainty for third order aberrations (Fig. 3).

**Figure 3.**
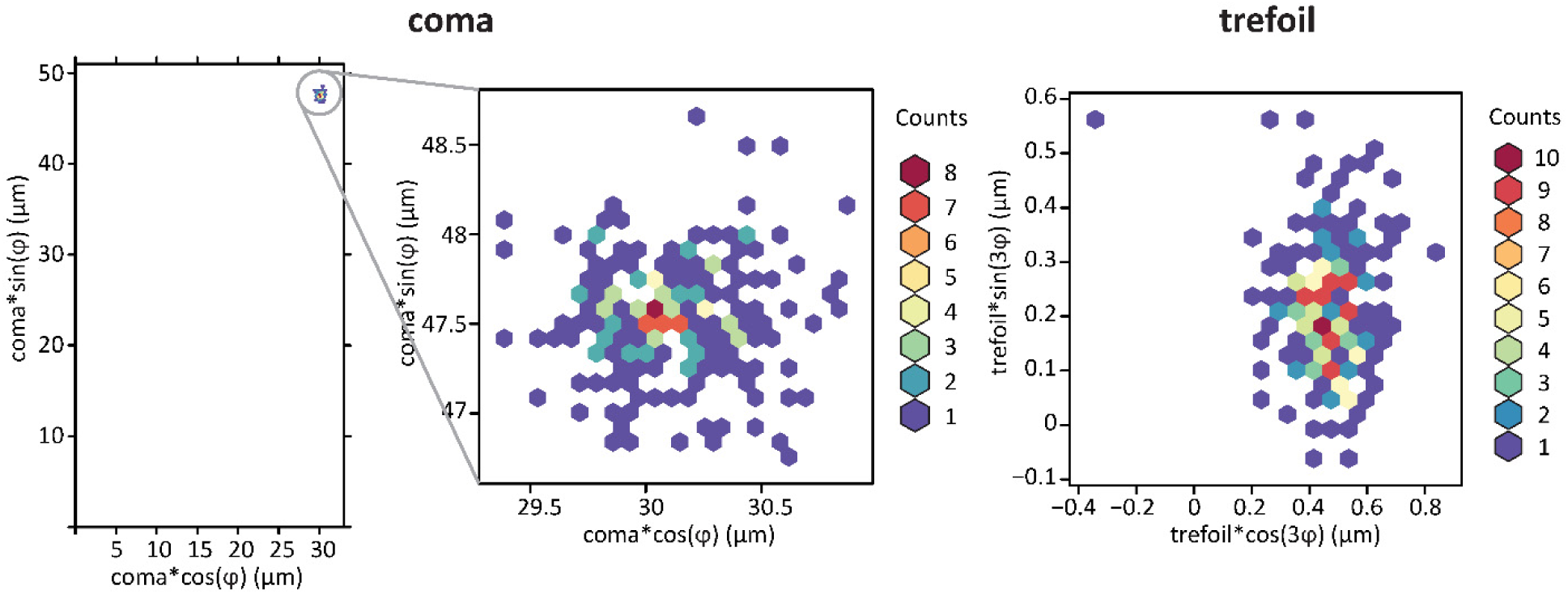
Heat maps for coma and trefoil values refined separately per image from the HemQ-57K dataset. The leftmost panel shows tight clustering of the coma values, with the center panel magnifying the region of clustering. The right panel shows the values of trefoil, which for this dataset were insignificant.

The problem generated by the presence of large coma was analyzed in a recent publication (11) which described the refinement of EMPIAR 10263, dataset III, with coma refinement performed by Relion 3.0 (13) and JSPR (11, 22), with results presented in Fig. 6B in (11). We reprocessed this dataset and obtained a much tighter clustering of coma values, similar to the clustering observed for our datasets. Relion 3.0 (13) seriously underestimated coma by a factor of ∼4 compared to our estimate and the average of the JSPR individual refinements per micrograph had a value that was underestimated by twenty-five percent compared to our values. However, we used a similar target function in our refinement procedure to JSPR (11, 22) and Relion 3.0 (13), and so the different outcomes are most probably due to differing refinement schemes (11). The results presented in Fig. 6B from reference (11) are consistent with the behavior of our procedure when compared to the results from our first cycle.

We observed that only the presence of a large fraction of incorrect particles or particles with grossly incorrect orientation in a micrograph would bias the coma refinement toward the starting point. When starting with coma refined by the previous cycle, re-refining the particle orientation attenuated the bias in the next cycle, so even the presence of a moderate number of bad particles did not affect the convergence of the procedure. We achieved additional improvement, in terms of resolution and spread of coma values, when we changed the null hypothesis regarding coma from a value of zero to the average of the refined coma values. This more appropriate null hypothesis was applied by taking advantage of the associative properties of coma, which allowed us to apply coma phase shift correction to the images; therefore, all the steps in the final round of the data analysis, starting from particle picking, were performed on images corrected by the average value of coma. Subsequent coma refinement, representing a difference from the previous average used in image correction, was still performed independently for all micrographs, but the residual bias resulting from the initial coma value being zero was eradicated. Coma or beam tilt refinement has to use the same type of reference-based target function irrespective of implementation. We would expect that small corrections would be equally well-characterized by all programs. However, local refinements and restraints may have different convergence ranges depending on the implementation, and this is mostly likely the explanation for the differences in results between programs.

Uncorrected aberrations create systematic patterns of phase shift and if such aberration error is constant across a dataset, then the reconstruction will also be altered in a systematic way, not only by losing amplitude but also by flipping its sign at some resolutions. In this case, the FSC curve may undergo oscillations, with the first minimum being the effective resolution limit of the result. Oscillations in the FSC curve are recognized as a qualitative sign of problems with the quality of cryo-EM SPR results (20). We provide here (Eq. 2) an explanation for another possible source of these oscillations. In our glucose isomerase (GI) data acquired with 58 μm coma, corresponding to 7.2 mrad beam tilt, before correcting coma we observed four oscillations in the FSC curve resulting from phase shift (Fig. 2) with the map interpretability being inconsistent with the FSC-based resolution indicator. Therefore, if FSC oscillations are encountered, refining coma is highly recommended to diagnose the problem, with the potential outcome being substantial reconstruction improvement by correcting the aberration.

## Discussion

Although instruments may be aligned quite accurately before data collection, our analyses and those of others (4, 6, 10, 11) indicate that coma can vary significantly between images and the extent of the variation is dataset dependent. For this reason, we recommend refinement of not only overall coma but also coma for individual images. Trefoil typically is not important, but for one pair of EMPIAR datasets (EMPIAR 10185 and EMPIAR 10186) and also our datasets (not shown), it was highly significant. However, even in these cases, only the overall value of trefoil for the dataset was important, with an insignificant level of variations between individual images.

Correction for antisymmetric aberrations can be performed during reconstruction (11, 13), but the consequence of the associative property is that it can just as well be performed at the whole micrograph level before reconstruction commences. In the presence of large coma, correcting coma at the image level reduces the point spread function of the imaging system so it may improve particle masking operations. If the value of these aberrations is known from external calibration, then the image-based correction may simplify applying these corrections to the data without modifying downstream analysis programs.

What affects cryo-EM SPRs is the error in the aberration model used and not the magnitude of the aberrations themselves, at least up to the theoretical limit where the image of a point source (e.g. coma) extends outside of the detector. Therefore, it is important to calibrate all aberrations and not only the ones that affect the power spectrum. This can be accomplished prior to data collection or a posteriori by reference-based refinement using the structure being solved. Once appropriate coma calibrations and corrections are used, this means that the procedures where the beam is intentionally tilted can be used on a larger scale. This can provide limited but additional three-dimensional particle information on top of the projection image without any time and precision penalties associated with mechanical rotations.

## Materials and methods

### Protein expression, purification, and grid preparation

Glucose isomerase (GI), also called xylose isomerase, from *Streptomyces rubiginosus* was purchased from Hampton Research (23). Protein slurry was dialyzed three times against excess of dH_2_O and concentrated to ∼40 mg/ml with Amicon filter.

HemQ protein was one of the structural genomics targets (MCSG APC35880). We have solved its X-ray crystallographic structure (PDB code: 1T0T.pdb) and recently others determined its function (24). Expression and purification of GYMC52_3505 plasmid encoding HemQ in pMCSG7 vector with Tobacco Etch Virus (TEV) cleavable N-terminal His_6_-tag (25) followed previously established protocol (26). After purification and tag cleaving, the protein was extensively dialyzed against 20 mM HEPES pH 7.5 and used at ∼28 mg/ml concentration for grid preparation. The plasmid GYMC52-3505 is available from the DNAS Plasmid Repository (https://dnasu.org). Cryo-EM grids for both proteins were prepared with FEI Vitrobot Mark IV. In each case, 3 µl of protein solution were applied to the grid at 4 °C, 100% humidity followed by 6 s blotting with blot force 20 before grids were plunged into liquid ethane cooled with liquid nitrogen (Fig S2).

### Data acquisition and analysis

The cryo-EM dataset for GI was collected with a 200 kV Talos Arctica microscope equipped with a K2 Gatan camera, with a physical pixel size of 0.91 Å. The phase plate was not used and the objective aperture was not inserted. A total of 202 movies with an exposure time of 100 s/movie were collected. Each movie contains 200 frames with an exposure time of 0.5 s/frame and an electron dose of 140 e/Å^2^ per movie (Table 1).

Both HemQ-57K and HemQ-45K were also collected in the same alignment conditions on a 200 kV Talos Arctica microscope with a K2 Gatan camera run in super-resolution mode, with a physical pixel of 0.72 Å for HemQ-57K and 0.91 Å for HemQ-45K. For HemQ-57K, 268 movies were collected with an exposure time of 40 s/movie. Each movie contains 100 frames with an exposure time of 0.4 s/frame and an electron dose of 90 e/Å^2^ per movie. For HemQ-45K, 257 movies were collected with an exposure time of 40 s/movie. Each movie contains 100 frames with an exposure time of 0.4 s/frame and an electron dose of 90 e/Å^2^ per movie.

Complete datasets for EMPIAR deposits 10204, 10185 and 10186 were processed as examples of datasets collected at 200 kV. EMPIAR 10185 and EMPIAR 10186 were collected consecutively on the same instrument with EMPIAR 10185 collected with a traditional setup by moving only the stage, and EMPIAR 10186 collected with the beam-image shift method. We performed image-specific correction for coma in both datasets, producing a material improvement in resolution (Table 2). EMPIAR 10263, dataset III, served as an example of a dataset collected at 300 kV with a large coma value that was partially corrected by alternative methods.

We processed all datasets with cisTEM (19). We modified the cisTEM pipeline by adding reference-based refinement of aberrations, including coma and trefoil, following the same design as implemented in Relion 3.0 (11, 13). As discussed in the Results and Discussion sections, multiple cycles that included aberration refinement, orientation refinement, and creation of a new reference (19). In these cycles, the resolution limit of data used for the orientation refinement was selected based on manual assessment of cisTEM’s SNR estimate exceeding a threshold value, typically around ∼4. The data collection and analysis statistics are summarized in Table 1. The cryo-EM movies and maps used in data analysis of GI and HemQ proteins are deposited in EMPIAR under [code 1] and [code 2] codes. For figure preparation, we placed models generated by crystallography (5VR0.pdb for GI and 1T0T.pdb for HemQ) into cryo-EM maps, using the rigid body refinement option available in Coot (27, 28).

## Acknowledgements

We thank Tabitha Emde for protein purification and grid preparation for the HemQ protein. We thank the Cryo-Electron Microscopy Facility at UT Southwestern Medical Center which is supported by grant RP170644 from the Cancer Prevention & Research Institute of Texas (CPRIT) for maintaining Talos Arctica microscope. This project has been funded in part with federal funds from the National Institute of Allergy and Infectious Diseases, National Institutes of Health, Department of Health and Human Services, under Contract No. HHSN272201700060C. This project was also supported by the National Institutes of Health (R21GM126406 to DB, and R01GM117080 and R01GM118619 to ZO) and the Department of Energy (DE-SC0019600 to YG).

## Author Contributions

RB, DB, and ZO developed an approach to aberration analysis; RB and ZO implemented the approach; DB acquired data; RB, YG and ZO analyzed data; RB, YG, DB and ZO wrote manuscript.

---

**Supplemental Figure S1.**
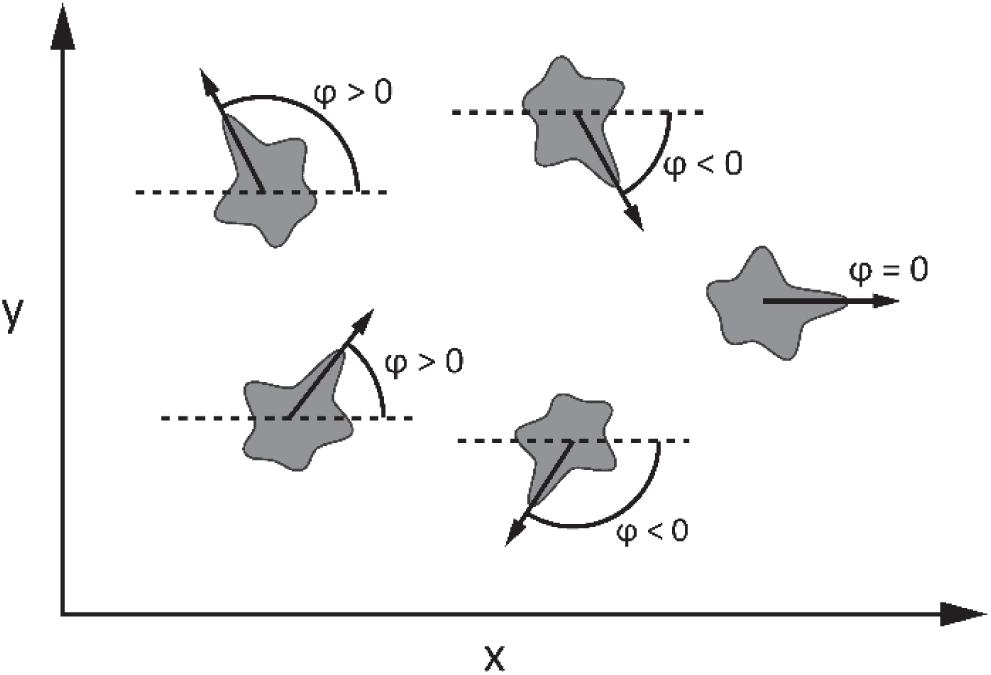
Definition of the orientation angle *φ* for particles coming from the same projection. This angle is not affected by forces generating preferred orientation in the typical setup of the sample grid being perpendicular to the beam.

